# Altering facial preferences with context

**DOI:** 10.64898/2026.05.27.727895

**Authors:** Asaf Madar, Naphtali Abudarham, Galit Yovel, Ido Tavor, Dino J Levy

## Abstract

Facial features are known to play a key role in how we judge a person’s traits. However, in many scenarios people do not evaluate a person in isolation but are required to compare and choose between multiple candidates. Yet, the principles that govern these decisions remain unknown. In economic literature, decisions are known to be influenced by the context of available options, but it is unclear to what extent these principles apply to social decisions about naturalistic stimuli such as human faces. Here, we bridged between these two fields of research - face evaluations and economic decisions - to investigate whether facial preferences can be systematically shifted by context, exhibiting a well-known choice context effect, the “decoy effect”. We asked participants to choose the person they think is more trustworthy from either a set of two (“target” and “competitor”) or three faces (including a “decoy”). Based on economic theory, we carefully selected face sets using their trustworthiness rankings and perceptual similarity, and tested them across several exploratory samples (n=100-400) and one large, preregistered experiment (n=1099). We found that adding a decoy face to the choice set increased participants’ choices of the target option by up to 7%, while the decoy face was rarely selected, consistent with a typical decoy effect. These results provide direct evidence that social judgements are affected by the same context-dependent principles known from economic choices, and that the decoy effect can be generalized to complex naturalistic stimuli such as human faces, relevant to various real-world applications.

**Significance statement:** People often rely on facial judgements when choosing between candidates, like politicians or potential partners. These judgments are typically made in comparison to available alternatives rather than in isolation. In economic literature, decisions are known to depend on the context of alternatives, but it remains unclear whether social evaluations follow the same principles. Here, tested whether the “decoy effect”, a well-known choice context effect, applies to social judgments. Participants were asked to choose the more trustworthy individual from either two (decoy-absent) or three faces (decoy-present). Adding a decoy face increased selection of the target face by up to 7%, while the decoy itself was rarely chosen, showing that social judgments are governed by the same context-dependent principles as economic choices.

## 1 Introduction

Humans make judgments about others all the time, deciding whom to trust, vote, approach or avoid, often based only on minimal visual information. Accordingly, these judgments are formed very quickly (Willis & Todorov, 2006) and show high agreement across individuals (Zebrowitz, 2011). Ample research identified several facial features related to these judgements (Todorov et al., 2015; Zebrowitz, 2017), as well as additional contextual cues such as the background or environment, that influence them (Brambilla et al., 2018; Geiselman et al., 1984; Mattavelli et al., 2023; Walker & Vul, 2014; Wang et al., 2020; Wu & Ying, 2023).

Notably, face judgements also have important real-world consequences, being associated with election outcomes (Olivola & Todorov, 2010; Todorov et al., 2005), financial performance (Graham et al., 2017; Rule & Ambady, 2008), and dating preferences (Little et al., 2006). Many of these scenarios involve a choice between multiple individuals, such as comparing politicians, job candidates, or potential partners. However, the principles that determine these choices are unknown. Interestingly, these choice situations closely resemble settings usually considered in economic frameworks, which analyzed multi-alternative choices and described plenty of context-dependent choice effects (Payne, 1982; Spektor et al., 2021). However, it is unclear to what extent the contextual principles known from the economic frameworks apply to the naturalistic settings of social judgments, when choosing between complex stimuli like human faces.

One of the most well-known context effects in economic choice is the *decoy effect* (Huber et al., 1982). The effect occurs when adding a third inferior “decoy” option to a set of two options, and while this third option is rarely selected, it still changes the tendency of choosing one of the original options (“target”) over the other (“competitor”). For example, cinemas could, and in many cases do, add a medium-sized high-priced bowl of popcorn to a menu already including a large high-priced bowl and a small low-priced bowl, to increase the sales of the large bowl (target) over the small one (competitor). Importantly, decoy options need to be carefully constructed in order to be effective, as they need to be both *inferior* and *similar* to the target (Huber et al., 1982, 2014; Simonson, 2014). The effect has been demonstrated many times since it was first described, but often only while using simple stimuli with explicit, two-dimensional numeric attributes, such as price and quality, where similarity and dominance relations are very easy to identify (Castillo, 2020; Choplin & Hummel, 2005; Dumbalska et al., 2020; Evangelidis et al., 2018; Gluth et al., 2017, 2018; Herne, 1999; Pettibone & Wedell, 2007; Simonson, 1989). Recent studies trying to use more naturalistic stimuli such as images of food products or movie banners, without explicit attributes and a structured presentation, have failed to find reliable decoy effects (Frederick et al., 2014; Trendl et al., 2021; Yang & Lynn, 2014), raising questions regarding whether the effect depends on artificial stimuli structures, which are rarely encountered outside the lab.

Face stimuli provide an excellent test for this question. Human faces are complex and naturalistic visual objects, lacking explicit numeric attributes, but elicit quick and reliable evaluations (Todorov et al., 2008), making them easy to use in a choice task with a natural setting. Here, we explored whether we could construct face stimuli sets that elicit the decoy effect known from economic choices. In other words, we tested whether the preference for a given face changes as a function of the identity of other faces which are present in the choice set and are available for choice. If choices between human faces exhibit the well-known context effects, the implications are twofold. First, these effects could provide concrete evidence for the generalization of the decoy effect to naturalistic stimuli and help develop economic theory that explains context effects in high-dimensional feature spaces. Second, real-world applications could use these effects to nudge people’s choices between potential candidates in a systematic, theory-driven way (e.g., dating apps, voting, etc.).

To test this, we built a pipeline to detect potential face triads that could elicit the decoy effect. First, we ranked faces by their perceived trustworthiness, focusing on this trait due to its reliability and quick evaluation process (Oosterhof & Todorov, 2008; Todorov, 2008; Willis & Todorov, 2006). Based on these rankings, we defined the *inferiority* between each pair of faces in our dataset (i.e., which face is more trustworthy). Second, we calculated the perceptual *similarity* between each pair of faces using explicit facial features and neural network representations. Consequently, we defined a few faces as targets and systematically searched for faces that potentially could serve as decoy and competitor faces. For the decoys, we searched for faces that are inferior to the target (less trustworthy), but at the same time perceptually similar to it, and for the competitors, we searched for faces equally trustworthy to the target, but perceptually different than it.

Finally, to test for a decoy effect, we asked participants to choose who they think is the most trustworthy person from a set of faces. Participants were assigned to one of two groups: a binary group, choosing between two faces (target and competitor), or a trinary group, choosing between the same two faces, but with an additional third face (decoy). The decoy effect was calculated as the difference in the propensity of choosing the target face between the two groups. In several exploratory studies and one large, preregistered experiment, we show that the presence of a decoy face reliably shifts preference toward the target, despite the decoy itself being rarely chosen. This provides direct evidence for a robust decoy effect using naturalistic stimuli and indicates that preference in social judgments is affected by the same contextual principles known from economic choices.

## 2 Results

In order to identify triads of faces that elicit the decoy effect on participants’ evaluations of face trustworthiness, we used a dataset of 100 male Caucasian faces. To reduce the heterogeneity in the stimuli, all faces were male Caucasian adults which had a neutral expression, no glasses, and no facial hair. Each face was also described by 20 numerical facial features such as skin color and jaw width, that were ranked previously by human observers (Abudarham & Yovel, 2016). Notably, one of the main challenges in finding such triads of faces, if exist, is the large number of unique ways to construct triads out of 100 faces (970,200 combinations). To narrow this enormous search space, we created a pipeline that identified triads that theoretically have high potential to elicit the decoy effect, by using two known properties that induce effective decoy options: inferiority and similarity to the target.

### 2.1 Estimating inferiority based on face trustworthiness

One important property of an effective decoy option is its inferiority relative to the target (Huber et al., 1982; Król & Król, 2019). Therefore, in order to find triads that elicit the decoy effect in trustworthiness evaluations, we started by estimating the difference in trustworthiness levels between faces, to find which faces are relatively inferior to other faces in this trait.

To do so, we collected responses from a sample of 126 participants who ranked the trustworthiness level of each of 100 faces that were used as the main dataset (Fig 1a-b). Overall, participants’ average rankings were highly reliable (Cronbach’s α=0.98, average fixed raters ICC=0.96), in line with previous reports (Todorov et al., 2008, 2009). The individual inter-rater agreement was relatively low (single fixed raters ICC=0.19), as reported in more recent studies analyzing the idiosyncrasies of subjective evaluations of faces (Martinez et al., 2020; Todorov et al., 2025). To define the level of inferiority between faces, we used the average normalized trustworthiness rankings across participants and calculated the difference between each pair of faces. We then used these differences as one of the two properties for constructing the face triads (Fig 1c).

**Figure 1.**
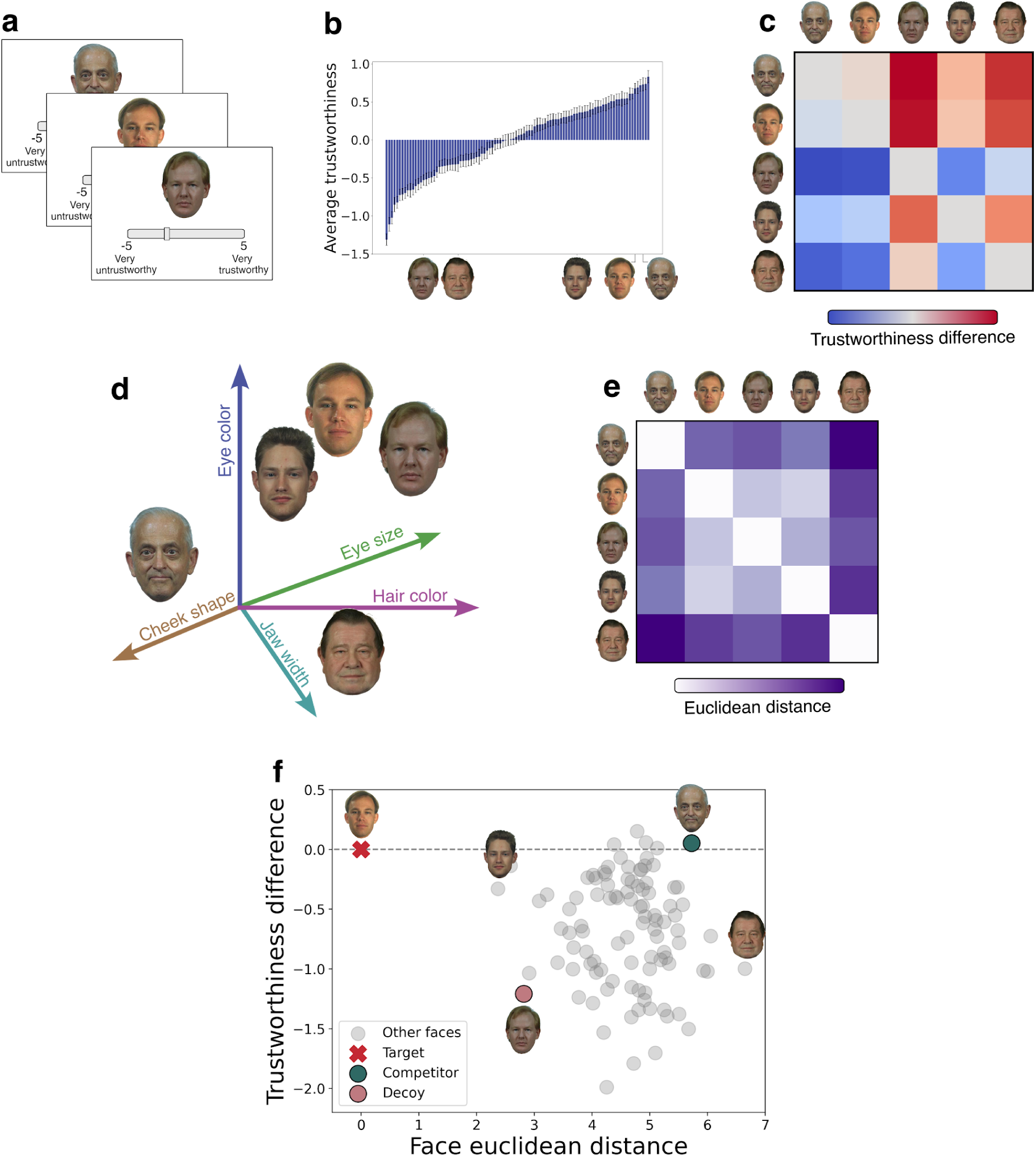
Pipeline for constructing face triads. (a-c) Face trustworthiness analysis. **(a)** One sample of participants (*n* = 126) ranked the trustworthiness of each face from a dataset of 100 faces. **(b)** The averaged normalized trustworthiness scores for each face across participants. Five example faces are marked on the X-axis. **(c)** We calculated the trustworthiness difference between each pair of faces. A small 5x5 difference matrix between the five example faces is shown. **(d-e)** Face representation similarity analysis. **(d)** Each face was represented on a twenty-dimensional space of facial features critical for face identity, as estimated by human observers in a previous study (Abudarham & Yovel, 2016). **(e)** We calculated the Euclidean distance between each pair of faces based on their positions in the face space. **(f)** We constructed triad of faces based on their distances in the face space (X axis, from panel e) and in trustworthiness (Y axis, panel c). The target is marked in red, competitor in green, decoy in pink, and other faces are marked in gray.

We also collected responses from another sample of only young adult males ( *n* = 201) to test whether reducing the heterogeneity of the sample increases the reliability of the trustworthiness rankings. First, the average rankings were highly similar between the samples (*r* = 0. 91). Surprisingly, the reliability results were also similar to the first sample, with high group-level reliability and low individual inter-rater agreement (Cronbach’s α=0.96, average fixed raters ICC=0.97, single fixed rater ICC=0.14). Nonetheless, we found that the young-male rankings were more tightly related to the faces’ facial features, as we were able to successfully predict this sample’s rankings, but not the heterogeneous sample’s rankings, using the facial features (Supplementary Materials 1).

### 2.2 Assessing the similarity between faces

The desired effect of a decoy option is that although it is not chosen often, it still affects the tendency to choose the target option in a non-trivial way. Importantly, simply adding a third face, inferior in trustworthiness levels, to the choice set would usually not result in shifting participants’ preferences towards any of the other options. Thus, to create the decoy effect, the decoy usually needs to be not just inferior, but also *similar* to the target (Huber et al., 1982; Tversky, 1977).

Thus, after defining the trustworthiness inferiority between each pair of faces in our dataset, we aimed to estimate their perceptual similarity. First, we represented the faces using an explicit facial feature space, including 20 facial features (e.g., skin color, jaw width, eye size, etc.; Abudarham & Yovel, 2016). Using these facial features, we calculated the dissimilarity between all 100 faces in our data set, and represent these values in a Representational Dissimilarity Matrix (RDM). Additionally, for robustness, as an alternative way to represent the faces we also used the faces’ representations in the penultimate layer of a DNN that was pretrained to classify face identity (Shoham et al., 2024), and calculated the DNN-RDM. The correlation between the two RDMs was *r* = 0. 30, suggesting the two feature spaces have a considerable overlap between their representations, but also that each contains unique information not included in the other.

### 2.3 Altering face preferences by adding a third decoy option

After assessing the trustworthiness difference and perceptual similarity between each pair of faces in the dataset, we constructed triads of faces that should theoretically elicit the decoy effect, based on these two properties (Fig 1f).

To do so, we first searched for target and decoy face pairs that show high perceptual similarity, while the decoy is inferior to the target in its trustworthiness level. Second, we searched for target and competitor pairs that show low perceptual similarity but have the same trustworthiness levels. Importantly, the target faces were arbitrarily defined and were not selected based on any specific parameters.

Based on these criteria, we constructed multiple face triads and tested if and to what extent they induce a decoy effect using several exploratory samples (*n* = 100 − 400, young male participants). Then, we selected the eight triads that showed the largest and most consistent decoy effects across all these samples and preregistered the results (https://aspredicted.org/gzvm-xhr3.pdf; Fig S1).

Next, to robustly test our results, we recruited a new large sample of male participants (*n* = 1099), which performed a face choice task using the eight preregistered triads. Participants were randomly assigned to either the binary (target and competitor faces) or trinary group (including a decoy face) and were asked to choose the most trustworthy face from the set (Fig 2a-b). We used the between-subjects design to better resemble real-world choice settings, which has also been commonly used in previous studies (Castillo, 2020; Frederick et al., 2014; Huber et al., 1982; Ratneshwar et al., 1987; Yang & Lynn, 2014). We calculated the size of the decoy effect for each face triad using the standard approach, as the difference in target choice ratios between the trinary and binary groups (Dumbalska et al., 2020; Gluth et al., 2018; Herne, 1999; Noguchi & Stewart, 2018; Pocheptsova et al., 2009; Rooderkerk et al., 2011; Spektor et al., 2018).

**Figure 2.**
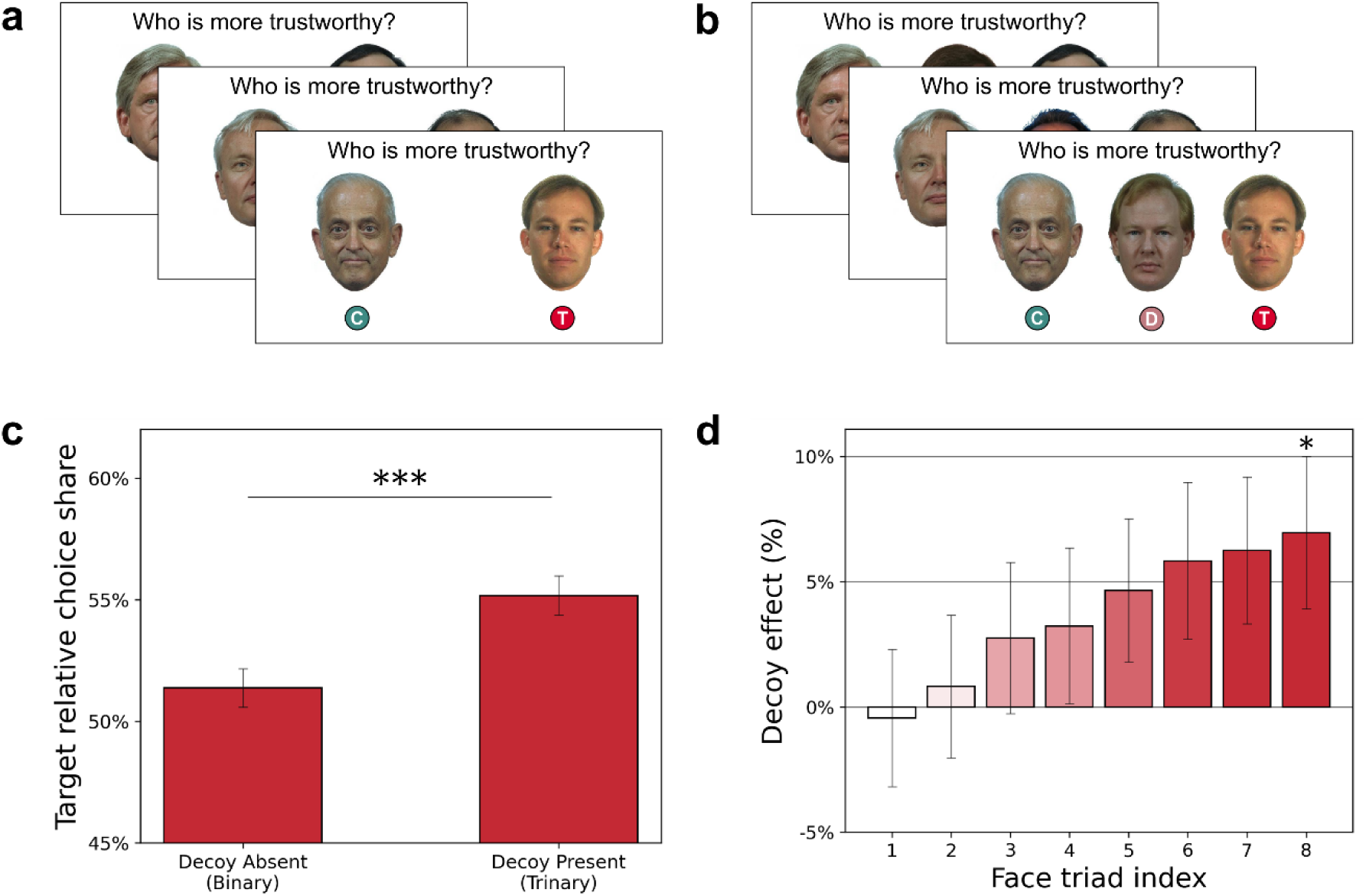
Face preferences change by the addition of a third decoy face. (a-b) The main choice task. Participants (*n* = 1099) were randomly assigned to either the binary **(a)** or trinary **(b)** group and asked to choose which face they think is more trustworthy. Participants in the binary group were presented with two faces (target and competitor), and participants in the trinary group were presented with the same two faces, with an additional decoy face. C – Competitor, D – Decoy, T – Target. **(c)** The propensity of participants who chose the target option in the binary and trinary groups, showing a significant decoy effect across all eight face triads. **(d)** The decoy effect for each face triad tested. Error bars represent bootstrap estimates of standard error. * *p* <0.05, *** , FDR corrected.

Across all eight face triads combined, adding a third decoy face to the choice set led participants to choose the target face as the most trustworthy 3.80% more compared to the same choice without a decoy present (55.17% vs. 51.37%; Fisher Exact Test, *p* = 0. 0003; Fig 2c). To also consider the dependency in choices within-participants, we fitted a mixed-effects logistic regression with random effects for participants and found a similar effect (β*_group_* = 0. 16, *p* = 0. 0007). Additionally, we tested whether repeated choices from three identical blocks of the same task exhibited the same effect and found similar results (56.28% vs. 52.84%; β*_group_* = 0. 15, *p* = 0. 0016). Importantly, participants chose the decoy face only rarely (range 7.29%-15.45%; mean 10.99%), meaning they perceived the inferiority in trustworthiness between the faces but were still affected by it, as intended. Moreover, as the target faces were arbitrarily defined, this shows we can potentially shift participants’ choices towards any face, given the right context.

As an exploratory analysis, we also calculated the decoy effect elicited by each face triad individually. As shown in Fig 2d, the triads elicited different magnitudes of effects, with 5 out of 8 showing effects larger than 3%. After testing their significance and correcting for multiple comparisons, only Triad 8 showed a significant decoy effect (*DE* = 6. 96%, Fisher Exact Test, *p* = 0. 0054) while Triad 6 and 7 did not survive the corrections ( *DE* = 6. 25%, *p* = 0. 0263; *DE* = 5. 83%, *p* = 0. 0337, respectively).

Together, our results show that choices based on social evaluations and complex stimuli exhibit the same contextual effects known from economic choices. Additionally, this provides direct evidence to the generalization of the decoy effect beyond structured stimuli and into naturalistic complex stimuli with no explicit numerical attributes, relevant to many real-world scenarios.

#### 2.3.1 Inferiority and similarity explain choices in the presence of the decoy

Next, we sought to explain what are the properties that lead participants to shift their preferences in the presence of the decoy option. We compared several known computational models of the decoy effect including value-normalization computational models (Adaptive Gain and Divisive Normalization), and logistic regression models using the faces’ similarity and inferiority values.

We trained the models to predict participants’ choices of the target option in the trinary group and evaluated them using a 5-fold cross-validation, such that different folds included different participants. For each model we calculated the average negative log likelihood (NLL) values on the held-out data. The computational models used the trustworthiness values of each face in the triad to predict choices, and the regression models used either the trustworthiness differences, the perceptual dissimilarity (based on the face space or DNN representations), or both, for each triad. The trustworthiness values were based on the rankings of the young-male ranking sample.

Comparing all models, we found that the regression model using inferiority and face space similarity reached the highest performance in predicting participants’ choices in the presence of the decoy option (*NLL* = 415. 1; Table S3). Additionally, the regression model using only inferiority performed better (*NLL* = 416. 8) than the one based only on face-space similarity (*NLL* = 423. 8), suggesting that although participants were affected by both the similarity and inferiority relationships between the faces, trustworthiness inferiority had a greater effect on participants’ choices. Finally, all regression models performed better than the best normalization-based model (Adaptive Gain, *NLL* = 548. 9), but note that the normalization-based models relied solely on trustworthiness, and not on the similarity information.

#### 2.3.2 Increasing the preference for the target face is driven by the decoy’s inferiority and dissimilarity to the target

Next, we analyzed the regressions’ coefficients to further explore what drove participants’ choices toward the target option to create the decoy effect.

First, as a baseline analysis, we trained an additional regression model to predict the choices of participants in the binary group, who chose without the decoy present, using the faces’ inferiority and similarity values. As expected, we found that participants chose the target more when the trustworthiness difference between the target and competitor increased (β = 0. 35), suggesting that they were sensitive to the trustworthiness rankings collected in the ranking sample. Also, as expected, the perceptual dissimilarity between the faces did not affect participants’ choices (β =− 0. 02). Finally, using the DNN representations to define dissimilarity showed very similar results ( Β*_trustwort_*_ℎ*iness*_ = 0. 36, β*_dissimilarity_*= 0. 04).

In the trinary group, participants’ choices were explained by a more complicated pattern of features (Table 1). First, like the binary group, participants’ target choices increased as the trustworthiness difference between the target and competitor increased ( β = 0. 59). Furthermore, we found that smaller trustworthiness differences between the target and decoy increased the probability to choose the target (β =− 0. 30), meaning that decoys that were closer to the target in their trustworthiness were more effective, in line with our hypothesis and previous works describing the importance of the decoy option’s inferiority property (Huber et al., 1982).

**Table 1.**
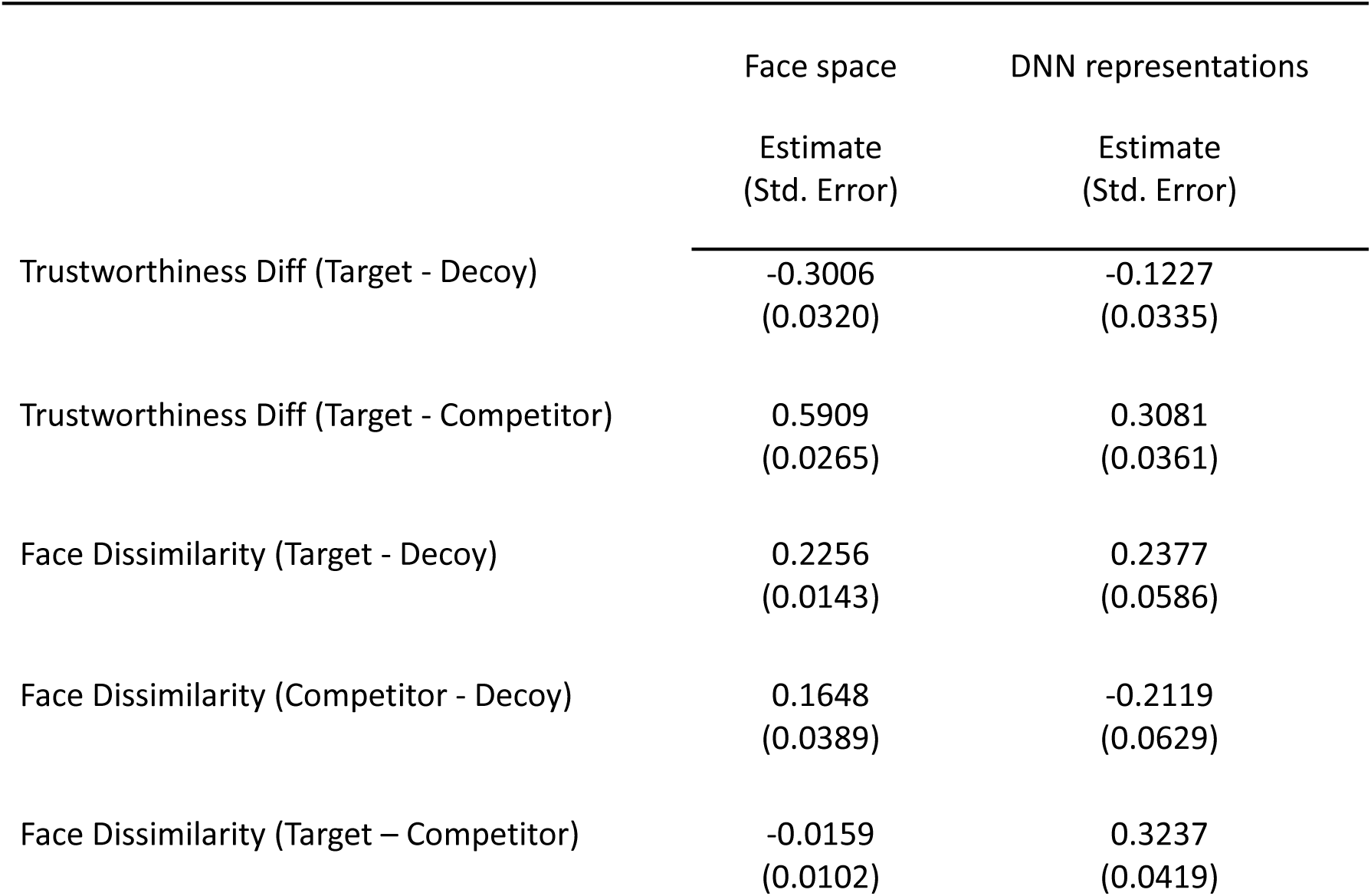
Coefficients of logistic regression models of the probability to choose the target option in the trinary group. Coefficients and standard errors were estimated using 5-fold cross-validation.

|  | Face space | DNN representations |
| --- | --- | --- |
|  | Estimate<br>(Std. Error) | Estimate<br>(Std. Error) |
| Trustworthiness Diff (Target - Decoy) | -0.3006<br>(0.0320) | -0.1227<br>(0.0335) |
| Trustworthiness Diff (Target - Competitor) | 0.5909<br>(0.0265) | 0.3081<br>(0.0361) |
| Face Dissimilarity (Target - Decoy) | 0.2256<br>(0.0143) | 0.2377<br>(0.0586) |
| Face Dissimilarity (Competitor - Decoy) | 0.1648<br>(0.0389) | -0.2119<br>(0.0629) |
| Face Dissimilarity (Target – Competitor) | -0.0159<br>(0.0102) | 0.3237<br>(0.0419) |

The perceptual dissimilarity between the target and decoy also played a role in participants’ choices. Namely, greater dissimilarity between the target and decoy in face space increased the probability to choose the target option (β = 0. 23), suggesting that the target-decoy similarity drove the decoy effect to the opposite direction than we intended. Greater dissimilarity between the competitor and decoy also affected the probability to choose the target option, but to a lesser extent (β = 0. 16). Finally, the dissimilarity between the target and competitor had no effect on participants’ probability to choose the target option, as intended (β =− 0. 02).

When repeating this analysis using the DNN representations instead of the face feature space to define perceptual dissimilarity, the results slightly differed but generally showed the same effects. First, like in the explicit face space model, greater dissimilarity between the target and decoy in DNN representations increased the probability to choose the target (β = 0. 24). On the other hand, greater target-competitor dissimilarity increased target choices (β = 0. 32), while the dissimilarity between the competitor and decoy decreased them (β =− 0. 21), effects that were not exhibited in the faces space model. Subsequently, the effects of trustworthiness differences were slightly weakened, but were still in the same direction as in the face space (target-decoy: β =− 0. 12; target-competitor: β = 0. 30). This suggests that the DNN representation distance may have encoded part of the faces’ trustworthiness difference too, resulting in the changed effects sizes compared to the face space.

For robustness, we also repeated these analyses by fitting the models to the entire dataset, instead of using cross-validation, and found very similar results (Supplementary Materials 2, Table S1, S2).

Together, these results suggest that choices of the target face were driven by the decoy’s trustworthiness inferiority to the target, as well as by their perceptual dissimilarity, as we hypothesized. Overall, participants were sensitive to the difference between the faces’ average trustworthiness rankings and were also affected by the trustworthiness values of the decoy option, despite rarely selecting it, exhibiting known economic choice context effects. Lastly, the perceptual similarity properties leading to an effective decoy were opposite than we hypothesized, as decoys more similar to the targets tended to *reduce* the probability to choose the target. This means that the pipeline to construct face triads could be improved by searching for target-decoy pairs that show greater dissimilarity, rather than the similarity patterns we aimed for.

#### 2.3.3 Stability of the face decoy effect

Finally, to estimate the stability of the decoy effect created by the face triads, we performed a bootstrap resampling analysis using the data from all exploratory and main samples (Fig 3).

**Figure 3.**
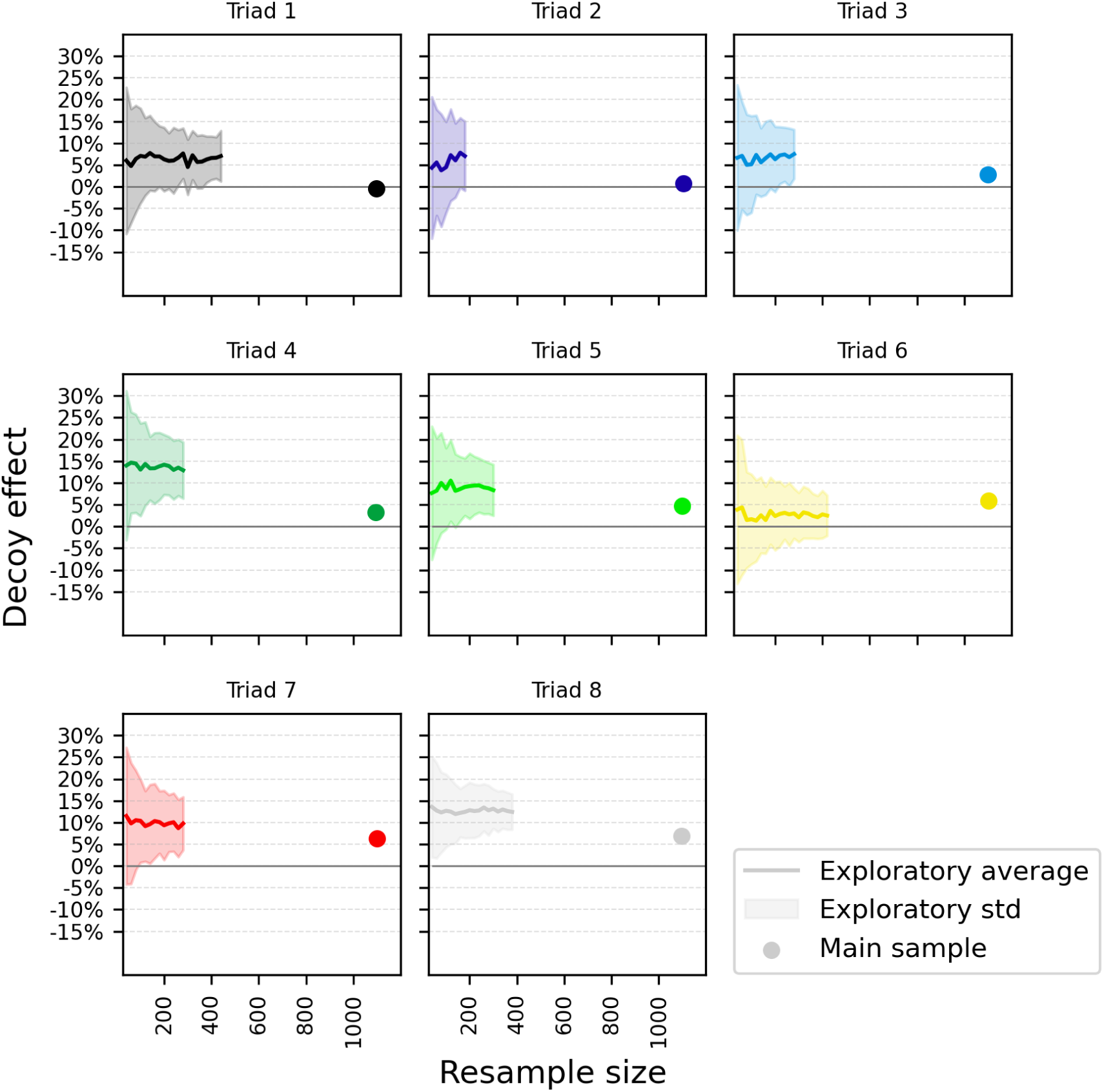
Stability of the decoy effect in the exploratory and main samples. For each face triad, we resampled the exploratory data with increasing sample sizes and calculated the average (line) and standard deviation (shaded area) of the decoy effect for each sample size. The decoy effects from the main sample are plotted as dots (*n* = 1099). The average distance between the main sample results and the exploratory average is -0.78 standard deviations.

This analysis had two aims. First, to estimate the variance of the decoy effect based on sample size, and second, to estimate the test-retest reliability for each face triad, comparing the effect calculated using the exploratory samples to the main sample.

To do so, we resampled the exploratory samples’ data 100 times with increasing sample sizes from *n* = 20 up to *n*≈400 (where possible, see Methods), and then calculated the average and standard deviation of the decoy effect for each sample size. As shown in Fig 3, all triads had a stable positive decoy effect, with standard deviations ranging from 4-8% (in the maximal sample size). Importantly, the results of the main sample were close to the results of the exploratory samples’ data, with an average distance of -0.78 standard deviations between them. This shows the effects are relatively stable across samples and sample sizes.

## 3 Discussion

In this work, we showed that choices based on social judgements exhibit systematic context effects, allowing us to shift people’s preferences toward an arbitrary face, given the right context. We presented participants with a set of either two (decoy-absent) or three (decoy-present) face stimuli and asked them to choose the face they think is more trustworthy. Using carefully selected sets of faces, we found a robust decoy effect, increasing participants’ choices of the target option by up to 7% when a third, inferior, decoy was present. Importantly, despite the decoy option being available for choice, it was only rarely selected, while still affecting participants’ choices of the other options.

To select these face sets, we relied on principles known from the traditional two-dimensional analysis of the decoy effect (Dumbalska et al., 2020; Huber et al., 1982; Król & Król, 2019) and applied them to face representations in high-dimensional feature spaces. Namely, we chose target-decoy face pairs that show perceptual similarity and trustworthiness inferiority, and target-competitor face pairs that show perceptual dissimilarity and equivalent trustworthiness levels. Investigating participants’ choices revealed that participants who chose in the presence of a decoy were affected by its inferiority but not by its similarity to the target face. These results demonstrate how complex social decisions could be manipulated using simple contextual cues in any direction, similarly to traditional economic choices.

The decoy effect in complex social choices is in line with considerable evidence for the effects of context on subjective evaluation of face traits. For example, negative and positive valence backgrounds have been shown to affect faces’ perceived trustworthiness (Brambilla et al., 2018; Wang et al., 2020; Wu & Ying, 2023). In another line of work, several studies showed how faces seem more attractive when surrounded by high attractive individuals than by low attractive ones (Carragher et al., 2019; Geiselman et al., 1984; Mattavelli et al., 2023; Walker & Vul, 2014). Nevertheless, these works focused on altering the perceived traits of a single face and have not tested the effect of context in multi-alternative choices, which is more relevant to daily social choices such as voting or dating. Collectively, these results show how the context, whether as a natural background or as surrounding individuals, can change the way we interpret crucial social cues from human faces and affect our choices.

Previously, Furl (2016) showed how choices between two faces based on their attractiveness level is affected by adding a third distractor face, an effect explained by divisive normalization models (Louie et al., 2013). Importantly, however, these context effects are unidimensional, based only on the faces’ attractiveness ratings, and can only reduce the number of participants choosing the more attractive face due to the introduction of the distractor. In contrast, the decoy effect, which can operate in a complex multi-dimensional space, can have an effect in both directions, either reducing or increasing the propensity to choose the target option. In our study, we selected the target option arbitrarily and aimed to increase its choice ratio, regardless of whether it was initially evaluated as a more trustworthy face. Model comparison results showed that the decoy effect in our study was explained better by trustworthiness inferiority and perceptual similarity than by the divisive normalization models which explained the distractor effect. Nonetheless, the divisive normalization models used in our work did not incorporate any similarity-related information, which could vastly improve their performance in this setting. Together, both the decoy and distractor effects strengthen the notion that complex stimuli such as human faces can create systematic context effects in choices. Future works could leverage insights from both types of effects to further develop computational models that fit complex stimuli such as human faces.

In the current study, we used a between-subjects design for the decoy choice task. In this design, each participant was presented with only one context condition, either with or without a decoy option, and the aim was to test whether the presence of the decoy changes the average preferences across participants. Notably, this approach resembles naturalistic choices, since people usually encounter only one type of choice set at a time in the real-world. Consequently, this design helps answer questions about the traits of the stimuli that create a choice bias on average at the population level for most people, rather than asking about the idiosyncratic choice of each individual, providing high relevance for real-world problems in marketing and choice architecture.

Interestingly, recent attempts using naturalistic stimuli to create the decoy effect have failed (Frederick et al., 2014; Trendl et al., 2021; Yang & Lynn, 2014). While the earlier attempts (Frederick et al., 2014; Yang & Lynn, 2014) ignited several methodological debates (Huber et al., 2014; Simonson, 2014), a more recent work generally resolved these issues but still found no effect (Trendl et al., 2021). Here, by carefully estimating the faces’ similarity, based on complex face feature spaces, and their inferiority, based on an independent ranking sample, we managed to find triads that consistently induce the decoy effect across large groups of participants. Additionally, using a between-subjects design helped us induce larger effects than the ones usually shown in previous within-subject studies and explore the stimuli’s properties that drive the effect. Thus, we show that the decoy effect can be extended to the domain of naturalistic stimuli such as faces, which are complex stimuli that were presented to participants without any explicit numerical attributes, giving direct experimental evidence for the generalization of the decoy effect to naturalistic complex stimuli.

Beyond its theoretical contribution, applying the decoy effect to face stimuli can have real-world consequences. As trustworthiness evaluations are known to be a general positive evaluation of faces (Oosterhof & Todorov, 2008), altering their perception could affect people’s decision whether to approach or avoid someone (Slepian et al., 2017; Todorov, 2008), whether in person or via websites and apps. Moreover, it is likely that a similar decoy effect could be found using other face traits, such as competence or attractiveness. Particularly, altering voters’ perception of candidates’ competence could potentially affect election outcomes (Todorov et al., 2005), and systematic biases in attractiveness could be relevant for decisions in dating apps.

The current experimental design has several limitations. First, while evaluations of trustworthiness are highly reliable at the group level, individual inter-rater agreement was much lower in our datasets, in line with previous reports (Martinez et al., 2020; Todorov et al., 2025). As the main choice task in this study was based on individual evaluations, it may be responsible for a large portion of the decoy effect’s instability exhibited in our samples. To control for this instability, we decreased the demographic heterogeneity of our sample and dataset, focusing only on male faces, and younger adult male participants. Future studies using a face trait with higher individual agreement could find even larger effects and in a more diverse sample of participants. Second, the final number of triads tested in this study was relatively low, partly due to the instability of the effects in the exploratory samples, which limited the ability to explore the facial features driving the effect. Future studies could increase the number of face triads and gain a better understanding of the mechanism underlying the decoy effect in this setting.

To conclude, our work shows that decisions over naturalistic social stimuli exhibit the decoy effect, by altering participants’ evaluations of face trustworthiness. These findings suggest that the decoy effect may extend to more ecological settings and could be applicable across a wide range of stimulus domains. When applied to faces, the decoy effect shows the role of contextual information in shaping social judgments which could have practical implications for real-world decision-making.

## 4 Methods

### 4.1 Face stimuli

The stimuli included 100 face stimuli from the FERET dataset (Phillips et al., 1998, 2000). The faces were male Caucasian adults which had a neutral expression, no glasses, and no facial hair. Their images were cropped from below the chin and up, as was used in previous works (Abudarham et al., 2019; Abudarham & Yovel, 2016).

### 4.2 Face trustworthiness ranking experiment

First, we aimed to estimate the faces’ trustworthiness rankings to define the level of inferiority between each pair of faces on the trustworthiness scale.

#### 4.2.1 Participants

We recruited an online sample of 145 participants. All participants were English speakers and lived in either the United States or the United Kingdom. Nineteen participants were excluded due to low variability in their responses (SD<0.5), extremely fast RT (mean RT<1s), or long duration for completing the task (>12 minutes). We thus analyzed the data of the remaining 126 participants. This sample was used for constructing the face triads.

An additional male-only sample of 232 participants was also recruited online via Prolific. All participants were male, English speakers, lived in either the United States or the United Kingdom, and their age was between 18-40 years. Thirty-one participants were excluded due to low variability in their responses, extremely fast RT, or if they failed at least one attention check. We thus analyzed the data of the remaining 201 participants. This sample was the last data collected in the experiment and therefore its rankings were not used for constructing the face triads, but only in the exploratory analyses.

#### 4.2.2 Ranking task procedure

Participants were presented with one face at a time and were asked to rate “How trustworthy is this person?” on a scale ranging from -5 (very untrustworthy) to 5 (very trustworthy, Fig 1a). We used the same instructions as was used in previous studies on trustworthiness evaluations (Adolphs et al., 1998; Todorov & and Duchaine, 2008). In detail, participants were informed the study was about first impressions and they were encouraged to imagine “trusting this person with all your money, or with your life, and then base your rating on this impression” and reacting based on their gut feelings. Participants rated all 100 faces in a random order and had no time limit for their responses. For the second sample, three additional attention checks appeared randomly throughout the task, showing a blank image (instead of a face), and asking participants to choose “5 (very trustworthy)”. The average time to complete the task was 8.8 min, and the average reaction time (RT) was 3.9s. Participants were paid an average of £1.11 for their participation.

#### 4.2.3 Ranking analysis

We normalized the rankings within-participant to zero mean and unit variance and averaged the normalized rankings across participants to reach an average trustworthiness ranking for each face stimulus (Fig 1b). To estimate the rankers’ agreement, we also calculated the Cronbach’s α and intra-class correlation (ICC) of the unnormalized rankings.

### 4.3 Face feature spaces and similarity scores

In order to construct face triads that would elicit the decoy effect, we assessed the perceptual similarity between the face stimuli, using two different feature spaces.

First, we represented each face by twenty explicit facial features (e.g., hair color, jaw width, etc.) that were evaluated by human observers in a previous work (Abudarham & Yovel, 2016). Each facial feature also had a weight corresponding to the inter-rater agreement between the raters who evaluated the features, where higher weight indicated higher agreement (Abudarham & Yovel, 2016).

Second, we represented each face using a convolutional deep neural network (DNN). We used a VGG-16 network (Simonyan & Zisserman, 2015) that was trained to classify face identity (Shoham et al., 2024), based on the VGGFace2 dataset (Cao et al., 2018). We first aligned the face stimuli using the MTCNN algorithm (Zhang et al., 2016) and then extracted the embeddings of the penultimate layer of the network for each face.

Finally, we calculated the distances between each pair of faces in each of the two attribute spaces and represented them in a 100 x 100 Representational Dissimilarity Matrix (RDM; Fig 1e). For the face feature space, we calculated a weighted Euclidean distance based on the features’ weights, and for the DNN representations we calculated the cosine distance.

### 4.4 Constructing face triads

To construct face triads that could potentially elicit the decoy effect, we applied the following logic, visualized in Fig 1f.

First, for the target and decoy options, we searched for possible decoy faces by identifying faces that have lower subjective trustworthiness evaluation compared to the target (inferior to the target), but at the same time perceptually similar to it. This means finding pairs of faces with a large trustworthiness difference but that are close in the perceptual feature space. Second, we searched for competitor faces that have similar subjective trustworthiness levels as the target face but perceptually different from it. We repeated this analysis for both types of perceptual similarity scores (face feature space and DNN representations).

To implement this logic, we first calculated the trustworthiness difference between each pair of faces, based on the subjective trustworthiness judgements of the ranking sample (Fig 1c). Then, we defined each of the 100 faces as the target option and searched for the best decoy face, i.e., the farthest in trustworthiness and closest in perceptual feature space. Then, we searched for the best competitor face, i.e., the closest in trustworthiness but the farthest in perceptual space. To find such pairs, we calculated two weighted scores for each pair of faces:

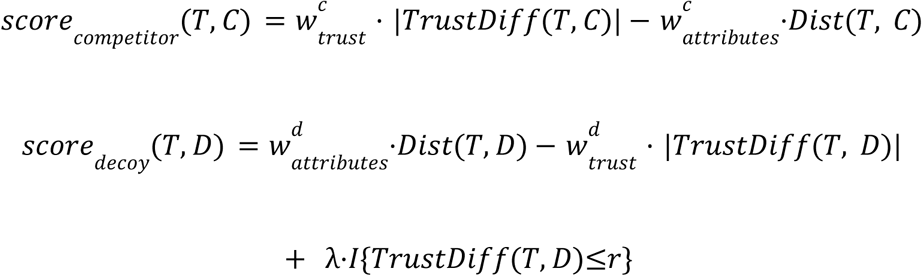

Where *T*, *C*, *D* are the target, competitor, and decoy faces, respectively, *TrustDiff* is the difference between the faces’ trustworthiness, *Dist* is the Euclidean or cosine distance between the faces in the attribute space, *w*∈*R* are independent weights for each score, and λ∈*R* is a dominance penalty for the decoy faces that promotes faces that are lower in trustworthiness compared to the target (in *r*∈*R* units or more).

The values of the free parameters λ, *w*, and *r* were chosen manually, based on visual inspection of the resulting face triads and their positions in Fig 1f (their trustworthiness difference and distance in perceptual space). To construct the final face triads, we selected the two faces which minimized *score_competitor_*, and *score_decoy_* to be the competitor and decoy faces, respectively. Finally, we dropped triads where the decoy was too close to the target in its trustworthiness and ensured each decoy face appeared only once across the triads. Overall, using variations of this pipeline we constructed 93 face triads out of the 970,200 possibilities to divide the 100 faces into unique triads.

#### 4.4.1 Filtering face triads

To find triads that create a robust decoy effect, we tested these 93 triads on several exploratory samples. Each sample viewed a different combination of triads, with each triad being tested by several hundred participants (ranging from 102 to 443, male participants aged 18-40). Based on the exploratory samples’ aggregated choices, we selected the eight face triads with the largest and most consistent decoy effects to be used as the final stimuli in the face choice task experiment (Fig S1). The exploratory results were also used for power analysis and preregistration before collecting the main sample (https://aspredicted.org/gzvm-xhr3.pdf).

### 4.5 Face choice task experiment

#### 4.5.1 Participants

We recruited 1300 participants online via Prolific. All participants were male, English speakers, lived in either the United States or the United Kingdom, and their age was between 18-40 years. The sample size was determined based on power analysis with π = 0. 8 and α = 0. 05. Two hundred and one participants were excluded due to one of the following: an average RT greater than 3 standard deviations from the mean RT of the exploratory samples, failing at least one attention check trial, or responding more than 90% of the time with the same key (e.g., left choice). We therefore analyzed the results of the remaining 1099 participants.

#### 4.5.2 Choice task procedure

Participants were randomly assigned to either the *binary* or *trinary* groups. In each trial, they were shown faces and were asked to choose which face they thought was more trustworthy. Participants in the binary group were presented with two faces in each trial. Participants in the trinary group were presented with three faces in each trial, which included the same two faces as the binary group with an additional decoy face. Each participant chose between eight unique face sets and performed three blocks of the task for a total of 24 trials. At the end of each block there was one attention check trial asking participants to ignore all previous instructions and answer a binary question: “Are you a human?”. After completing these three blocks, participants faced additional three attention trials, asking them to choose the face with the darkest hair, from a set of either two or three faces, in a similar design to the main task. Participants who failed in more than one attention check were excluded. The average time to complete the task was 3.5 min. Participants were paid £0.5 for their participation.

#### 4.5.3 Decoy effect analysis

To test whether the addition of the third decoy face affected the probability of participants to choose the target face, we calculated the between-subjects decoy effect for each face triad. First, we calculated the target relative choice ratio (RCS), for the trinary and binary groups:

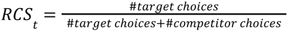

Where *t*∈{1, …, 8} is the face triad index. Following previous studies, the magnitude of the decoy effect (DE) was defined as the difference between the trinary and binary RCS (Dumbalska et al., 2020; Gluth et al., 2018; Herne, 1999; Noguchi & Stewart, 2018; Pocheptsova et al., 2009; Rooderkerk et al., 2011; Spektor et al., 2018), representing the relative increase in target choices attributed to the addition of the decoy option:

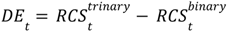

This resulted in eight decoy effects, one for each face triad in the choice task.

As preregistered, all choice ratios were calculated using only the first block of choices, in order to evaluate the effect on participants’ first impressions of the faces. We used the additional two blocks to examine the influence of repeated choice on the decoy effect in this setting. We tested the decoy effect significance using a one-sided Fisher Exact Test (expecting an increase in the targets’ choice ratios).

To account for the dependency in choices within-participants, we also fitted a mixed-effects logistic regression to explain the probability of choosing the target face with random effects for participants, a fixed effect for group (trinary/binary), and control variables including RT, age, and ethnicity. Control regressors were normalized to zero mean and unit variance before fitting. We tested whether the coefficient for the group regressor is different from zero using a one-sided t-test (expecting an increase in target choices in the trinary group).

For all samples and analyses, we dropped trials that were performed extremely fast (<50ms) and extremely slow (more than 5 standard deviations from the sample’s mean), affecting 0.34% of trials in the main sample.

#### 4.5.4 Computational modelling

To explain the potential driving factors for the decoy effect, we trained several computational models to predict participants’ choices in the trinary group, who chose in the presence of the decoy option. We trained the models to predict the choice of the target option and evaluated them using a 5-fold cross-validation, where the folds included different participants. We calculated the average negative log likelihood (NLL) values on the held-out participants’ choices and compared the models’ performance. All models were trained only on target and competitor choices, excluding a small portion of decoy choices.

##### 4.5.4.1 Value normalization models

First, we used previously suggested computational models which are based on value normalization, including Adaptive Gain (Dumbalska et al., 2020), Recurrent and standard Divisive Normalization (Dumbalska et al., 2020; Furl, 2016; Webb et al., 2020). These models applied normalization on the faces’ trustworthiness rankings to predict participants’ choices of the target option, and have been shown to predict the decoy effect in other stimuli types.

To train the models, we scaled the trustworthiness values of all faces to be between zero and one using min-max scaling. Then, we fitted the model’s parameters to the train participants’ choices using the Nelder-Mead simplex algorithm implemented by *scipy*’s *fmin* function (Virtanen et al., 2020), and applied the model to predict the test participants’ choices.

We used the Divisive Normalization model (Dumbalska et al., 2020; Webb et al., 2020), which estimates each option’s subjective value by its raw value divided by the sum of all options’ values:

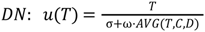

Where *u*(⋅) is the utility function, and *T*, *D*, and *C* are the trustworthiness values of the target, decoy, and competitor. σ and ω are free parameters representing the bias and weight of the normalization procedure.

Next, we used the Recurrent Divisive Normalization (Dumbalska et al., 2020; Webb et al., 2020), which is similar to the Divisive Normalization model, but overweighs the value of the estimated option:

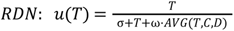

Finally, we used the Adaptive Gain model which assumes the utility of each choice option is dependent on the context via a sigmoidal nonlinearity:

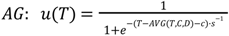

Where *s* is the sigmoid slope and *c* is the sigmoid bias (together with the average trustworthiness of *T*, *D*, *C*).

Finally, the choice was made by a softmax function with temperature parameter τ:

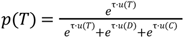

##### 4.5.4.2 Regression models

To test the effects of trustworthiness inferiority and perceptual similarity on participants’ choices, we trained several logistic regressions to predict the probability to choose the target face in each group. The regressors included the trustworthiness difference between each pair of the available options (binary group: target-competitor; trinary group:

target-competitor, target-decoy, and competitor-decoy), and the dissimilarity between each pair of available options. All regressors were normalized to zero mean and unit variance.

The simplest regressions included either the inferiority or similarity values, and the full regression models included both properties:

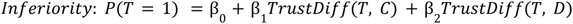

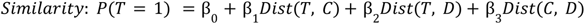

We used the trustworthiness values based on the rankings of the young-adult sample. We defined the dissimilarity by the face feature space in one type of models, and by the DNN representations in another type of models. In total, we trained five models: one inferiority model, two similarity models, and two full models.

Finally, for the models’ coefficients analysis, we calculated the average coefficient and standard deviation over the five folds of the cross-validation.

#### 4.5.5 Decoy effect stability analysis

To evaluate the stability of the decoy effects, we used resampling techniques on the exploratory results. Specifically, we resampled with replacement using increasing sample sizes (*n* = 40, 50, …, *N*) and calculated the decoy effect in each resampled data, repeating the procedure 100 times for each sample size. Then, we calculated the average and standard deviation of the decoy effect for each sample size. The maximal sample size for each triad was determined by the number of participants in the exploratory data that completed the task with this triad, as not all participants in the exploratory samples viewed the same sets of triads (min *N* = 177, max *N* = 443).

### 4.6 Data and code availability

All analyses codes and raw behavioral data are available at https://github.com/asafmm/face_decoy. Portions of the research in this paper use the FERET database of facial images collected under the FERET program, sponsored by the DOD Counterdrug Technology Development Program Office. You may not use any of the images in this manuscript without written permission from NIST and from the authors.

## 4.7 Acknowledgements

The authors acknowledge with thanks the support of the Israel Science Foundation (1432/23) and the Henry Crown Institute of Business Research in Israel.

## 4.8 Author contributions

A.M., N.A., G.Y., I.T., and D.J.L. designed the research. A.M. designed the experiments, collected the data, and performed the analyses. A.M., G.Y., I.T., and D.J.L. wrote and edited the paper.

