## Supplementary Materials for "Altering facial preferences with context"

### 1. Predicting face trustworthiness

As a secondary aim and exploratory analysis, we tested the ability to predict trustworthiness and explored the relationship between facial features and trustworthiness. We trained models to predict the trustworthiness of each face, based on the rankings of the young-male sample. To do so, we trained machine-learning (ML) and DNN models to predict the average trustworthiness rankings. The ML models were trained to predict trustworthiness using the faces' explicit facial features, and the DNN was trained to predict it directly from the images.

First, we trained a Lasso regression model to predict trustworthiness based on the facial features from the face space. The model was evaluated using leave-one-out cross-validation and the regularization parameter was tuned using nested 5-fold cross-validation. The values of regularization were set automatically by python's scikit-learn (Pedregosa et al., 2011).

Using the Lasso model, we significantly predicted the trustworthiness from the faces' facial features ($r = 0.48$, $p < 10^{-6}$; $MSE = 0.83$). Exploring the coefficients of the Lasso models revealed that lighter skin color, smoother skin texture, shorter forehead height, shorter hair length, and darker hair color predicted higher trustworthiness (features are ordered from the most to least important). Interestingly, when training the model to predict the trustworthiness rankings from the more heterogenous, first ranking sample, the predictions where much worse ($r = 0.18$, $p = 0.076$;$MSE = 1.00$), showing that although the inter-rater agreement metrics were pretty similar between the samples (see Section 2.1 in Main Text), the young male sample's average rankings were more coherent in their relationship to the facial features. Additionally, this effect was not related to the larger sample size of the young-male sample (n=201) compared to the heterogenous sample (n=126), as resampling n=126 without replacement from the young-male sample did not change the result (mean $r = 0.47$, $MSE = 0.84$ across N=100 resamples).

Additionally, we fine-tuned the VGG-16 face recognition network used for extracting the DNN representations (Shoham et al., 2024). We replaced the output layer to predict a single value and trained the network to predict trustworthiness directly from each image. All weights were unfrozen for the training phase. The network trained on 80 faces and tested on a left-out set of 20 faces. The hyperparameter tuning was based on a 5-fold nested-cross validation within the training set. The final training included 10 epochs, and the learning rate decreased by a factor of 10 when the loss reached a plateau on a validation set (10% of the training set). The trustworthiness scores were normalized based on the training data. The training and hyperparameter tuning were implemented using PyTorch and Ray Tune (Liaw et al., 2018; Paszke et al., 2019).

Using the fine-tuned DNN, we achieved a slightly worse performance than the Lasso model ($r = 0.38$, $MSE = 0.89$). Nonetheless, the advantage of the network over the linear model is that it is not limited to faces that have their explicit facial features labeled.

### 2. Fitting choices using trustworthiness inferiority and perceptual similarity

To test the effect of trustworthiness rankings and perceptual similarity on participants' choices, we fitted mixed-effects logistic regressions to explain the probability to choose the target face in each group. The regressors included fixed effects for the trustworthiness difference between each pair of the available options (binary group: target-competitor; trinary group: target-competitor, target-decoy, and competitor-decoy), and the dissimilarity between each pair of available options, a random effect for participants, and control variables of RT, age, and ethnicity. We fitted one regression using the dissimilarity based on the face feature space, and another one based on the DNN representations. All regressors were normalized to zero mean and unit variance before fitting.

In the binary group, participants' target choices increased as the trustworthiness difference between target and competitor increased ($\beta = 0.36$, $p < 10^{-15}$), suggesting that they were sensitive to the size of the average trustworthiness difference between the faces. Also, as expected, the perceptual dissimilarity between the faces did not affect their choices ($\beta = -0.03$, $p = 0.36$). Lastly, using the DNN representations instead of the face feature space for perceptual dissimilarity yielded very similar results (see Table S1).

**Table S1.** Estimates of a mixed-effects logistic model of the probability to choose the target option in the binary group.

| | Face space | DNN representations |
|---|---|---|
| | Estimate (Std. Error) | Estimate (Std. Error) |
| Intercept | 0.0630 (0.0350) | 0.0620 (0.0350) |
| Trustworthiness Diff (Target - Competitor) | 0.3565 *** (0.0341) | 0.3702 *** (0.0332) |
| Face Dissimilarity (Target - Competitor) | -0.0293 (0.0323) | 0.0466 (0.0314) |
| Age | 0.0364 (0.0356) | 0.0365 (0.0356) |
| White Ethnicity | -0.0373 (0.0357) | -0.0373 (0.0357) |
| Reaction Time | -0.1018 ** (0.0336) | -0.1005 ** (0.0335) |
| Log Likelihood | -2947.0 | -2946.3 |
| Individuals | 547 | |
| Observations | 4366 | |

*Note*. The model also includes random effects at the participant level. All regressors were normalized before fitting. * $p < 0.05$, ** $p < 0.01$, *** $p < 0.001$.

In the trinary group, participants' choices were explained by a more complicated pattern of features (Table S2). First, like the binary group, participants' target choices increased as the trustworthiness difference between the target and competitor increased ($\beta = 0.57$, $p < 10^{-10}$). Moreover, smaller trustworthiness differences between the target and decoy increased the probability to choose the target ($\beta = -0.28$, $p = 0.0068$), meaning that decoys that were closer to the target in their trustworthiness were more effective, in line with our hypothesis and previous works of the decoy effect (Huber et al., 1982). The perceptual dissimilarity between the target and decoy also played a role in participants' choices. Namely, greater dissimilarity between the target and decoy in face feature space increased the probability to choose the target option ($\beta = 0.24, p < 0.0001$), suggesting that the target-decoy similarity drove the decoy effect to the opposite direction than we intended.

When repeating this analysis using the DNN representations instead of the face feature space to define dissimilarity, the results slightly changed but generally showed the same effects. First, like in the face space analysis, greater dissimilarity between the target and decoy in DNN representations increased the probability to choose the target ($\beta = 0.28$, $p = 0.0025$). On the other hand, greater target-competitor dissimilarity increased target choices ($\beta = 0.35$, $p = 0.0005$), while the dissimilarity between the competitor and decoy decreased them ($\beta = -0.24$, $p = 0.0037$), effects that were not exhibited in the faces space analysis. Subsequently, the effect of trustworthiness difference was slightly less significant, but still in the same direction as in the face space (target-decoy: $\beta = -0.12$, $p = 0.0564$; target-competitor: $\beta = 0.29$, $p = 0.0003$). This suggests that the DNN representation distance may have encoded part of the faces' trustworthiness difference too, resulting in the changed effects sizes compared to the face space. Nonetheless, as the DNN-based model had an overall worse fit than the face space model, we focus on the face space model interpretation ($NLL_{DNN} = 2589.8$, $NLL_{Face\ space} = 2583.0$; see Table S2).

**Table S2.** Estimates of a mixed-effects logistic model of the probability to choose the target option in the trinary group.

| | Face space | DNN representations |
|---|---|---|
| | Estimate (Std. Error) | Estimate (Std. Error) |
| Intercept | 0.2403 *** (0.0399) | 0.2319 *** (0.0396) |
| Trustworthiness Diff (Target - Decoy) | -0.2801 ** (0.1035) | -0.1248 (0.0654) |
| Trustworthiness Diff (Target - Competitor) | 0.5741 *** (0.0871) | 0.2877 *** (0.0804) |
| Face Dissimilarity (Target - Decoy) | 0.2401 *** (0.0577) | 0.2812 ** (0.0928) |
| Face Dissimilarity (Competitor - Decoy) | 0.1405 (0.0771) | -0.2413 ** (0.0832) |
| Face Dissimilarity (Target - Competitor) | -0.0068 (0.0684) | 0.3500 *** (0.0999) |
| Age | 0.0008 (0.0402) | 0.0016 (0.0401) |
| White Ethnicity | -0.0548 (0.0403) | -0.0527 (0.0402) |
| Reaction Time | -0.1859 *** (0.0373) | -0.1786 *** (0.0371) |
| Negative Log Likelihood | 2583.0 | 2589.8 |
| Individuals | 552 | |
| Observations | 3913 | |

*Note*. The model also includes random effects at the participant level. All regressors were normalized before fitting. The feature "Trustworthiness Diff (Comeptitor-Decoy)" was omitted from the regression as it is a linear combination of the two other trustworthiness features. * $p < 0.05$, ** $p < 0.01$, *** $p < 0.001$.

**Table S3.** Model performance for computational and regression models predicting participants' choices in the trinary group. Negative log likelihood (NLL) values show the average NLL on held-out data in a 5-fold cross-validation procedure.

| **Model Type** | **NLL** |
|---|---|
| Divisive Normalization | 619.5 |
| Recurrent Divisive Normalization | 595.7 |
| Adaptive Gain | 548.9 |
| Face-space Similarity | 423.8 |
| DNN Similarity | 417.2 |
| Inferiority | 416.8 |
| Inferiority & DNN Similarity | 416.3 |
| **Inferiority & Face-space Similarity** | **415.1** |

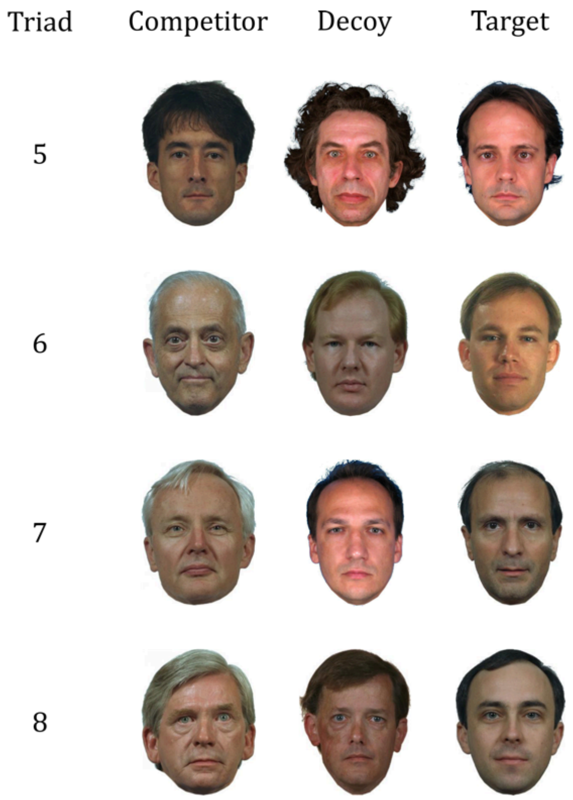


**Figure S1. Top face triads used for the main sample.** These face triads were constructed based on different configurations of the pipeline detailed in Fig 1, and showed the largest and most stable effects in the exploratory data. Triads are ordered by the decoy effect they elicited in the main sample (Fig 2). To comply with the FERET dataset terms of use, only a subset of the face images is presented here.
